# Neuro-toxicogenomic mapping of TMT induced neurotoxicity using human minibrain reveals associated adverse outcome events

**DOI:** 10.1101/2022.01.21.477206

**Authors:** Govindan Subashika, Alpern Daniel, Luc Stoppini, Bart Deplancke, Roux Adrien

## Abstract

Genome-wide transcriptomic interrogation of organoids and 3D tissue models are increasingly used for characterizing drug, toxicity responses and neurodevelopmental disorders. We established here a neuro-toxicogenomic assay by utilizing “minibrain”, a human *in vitro* 3D brain model system and a low-cost, highly multiplexable RNA-seq methodology (BRB-seq) for screening the effect of trimethyltin chloride (TMT) induced neurotoxicity. We demonstrate that transcriptomic profiling is insightful to the cellular composition and regional identity of the minibrain. Further, we characterize the transcriptomic changes associated with the dose-time neurotoxic response of minibrain upon exposure to TMT. The distinct gene expression changes and molecular candidates identified with our pipeline provides insight to map the key events involved in the adverse outcome pathway of TMT associated neurotoxicity. We identify processes such as endoplasmic reticulum stress, dysregulation of synaptic genes and downregulation of neuron-morphology associated genes upon exposure to TMT. In response to TMT, we identify activation of an early response homeostatic mechanism in minibrain and an interplay of STAT pathways correlating with the dose severity. In this study, we present a neuro-toxicogenomic assay that demonstrates the power of a low-cost transcriptomic screening to study chemical induced neurotoxicity.

## 1 Introduction

Routine exposure to chemicals has increased in humans in modern times causing toxic effects on both the ecosystem and human body. Neurotoxic effects of many of these chemical compounds on behavior have been established in different experimental models. Among these chemicals, organotins are one of the endocrine-disrupting chemicals (EDC) with potent neurotoxic actions on the central nervous system (Ferraz da Silva *et al*., 2018; Sandström *et al*., 2019). Trimethyltin chloride (TMT) is one of the most common organotin class toxins used as fungicide and plastic stabilizers known for its neurotoxic effects in humans (Kreyberg et al., 1992; Wang et al., 2017). In mice, TMT has been shown to cause dysregulated firing of neurons causing seizures, neuronal death and reduction in glutathione homeostasis causing neuronal insult (Shin *et al*., 2005).

The advent of *in vitro* 3D brain models and low-cost transcriptomics now allow us to study the effects of toxicants on human neurons and brain microenvironment at the molecular level in real time (Zander et al., 2017; Brown et al., 2018; Liu et al., 2019; Sandström et al., 2019). In this study we use minibrains developed in our lab as an in *vitro* model to map molecular events that lead to TMT induced neurotoxicity. Previously we described a pipeline to mass generate human minibrains, perform large scale immunolabeling, viral mediated neuron labeling and 3D imaging that would facilitate reconstruction of neuron morphology and detection of molecular markers in 3D for screening studies (Govindan et al., 2021). Here, we utilize BRB-seq, an ultra-low-cost transcriptomics pipeline that allows time and cost-efficient multiplexing of large numbers of samples to setup our neuro-toxicogenomic assay in exposure to TMT. Using the neuro-toxicogenomic assay we identified the dose-time TMT exposure induced transcriptional response in minibrains (Figure 2). The neuro-toxicogenomic assay allowed us to map biological events that dynamically occur across exposure timepoints and toxicity severity of TMT (Figure 2–5).

## Method

### Minibrain generation and TMT neurotoxicity assay

Neural stem cells derived from induced pluripotent stem cells (NSC^hIPSC^) (ThermoFisher, #A3890101) were seeded with 2E5 cells per 6 well-plates and processed for minibrain generation according to the protocol previously described (Govindan et al., 2021). Trimethyltin chloride (TMT) (Merck-Sigma 146498) compound at a concentration of 0,1, 2.5, 5 μM was added to 4-month-old mature minibrains and was treated for 1 day (acute), 3 days (intermediate) and 7 days (chronic).

### Immunohistochemistry and neuron labeling in minibrain

Minibrains were fixed with 4% paraformaldehyde and stained with anti-GFAP (Invitrogen, 180063), anti-beta TUBIII (Sigma, T8660) and anti-SYNAPSIN (sigma, SAB4502903) as described in (Govindan et al., 2021). Neurons in the minibrain were labelled with tdTomato through transduction of retrograde viral particle carrying TdTomato reporter (AAVRg:tdTomato) as described in (Govindan et al., 2021) Minibrains were mounted in RapiClear 1.47 and imaged using a confocal microscope.

### RNA isolation and BRB-seq library preparation

For characterizing the minibrain transcriptome in Figure 1, 1 million NSC^hIPSC^ (3 replicates) and 3-month-old minibrains (6 replicates) were harvested. For the neuro-toxicogenomic assay 4-month-old TMT treated minibrains were harvested across all treatment conditions (3 technical replicate/ condition). RNA was extracted from the samples according to the Qiagen protocol (Qiagen, RNA easy plus kit, 74034). 10 μl of RNA eluate corresponding to 50-100 ng/μl was transferred into 96-wellplate (low bind, RNase-free, Eppendorf, 0030129504). The generation of bulk RNA Barcoding and sequencing (BRB-seq) libraries was performed using MERCURIUS BRB-seq kit and following manufacturer manual (Alithea Genomics, #M1001), which represents an improved version of the original BRB-seq protocol (Alpern et al., 2019).

**Figure 1:**
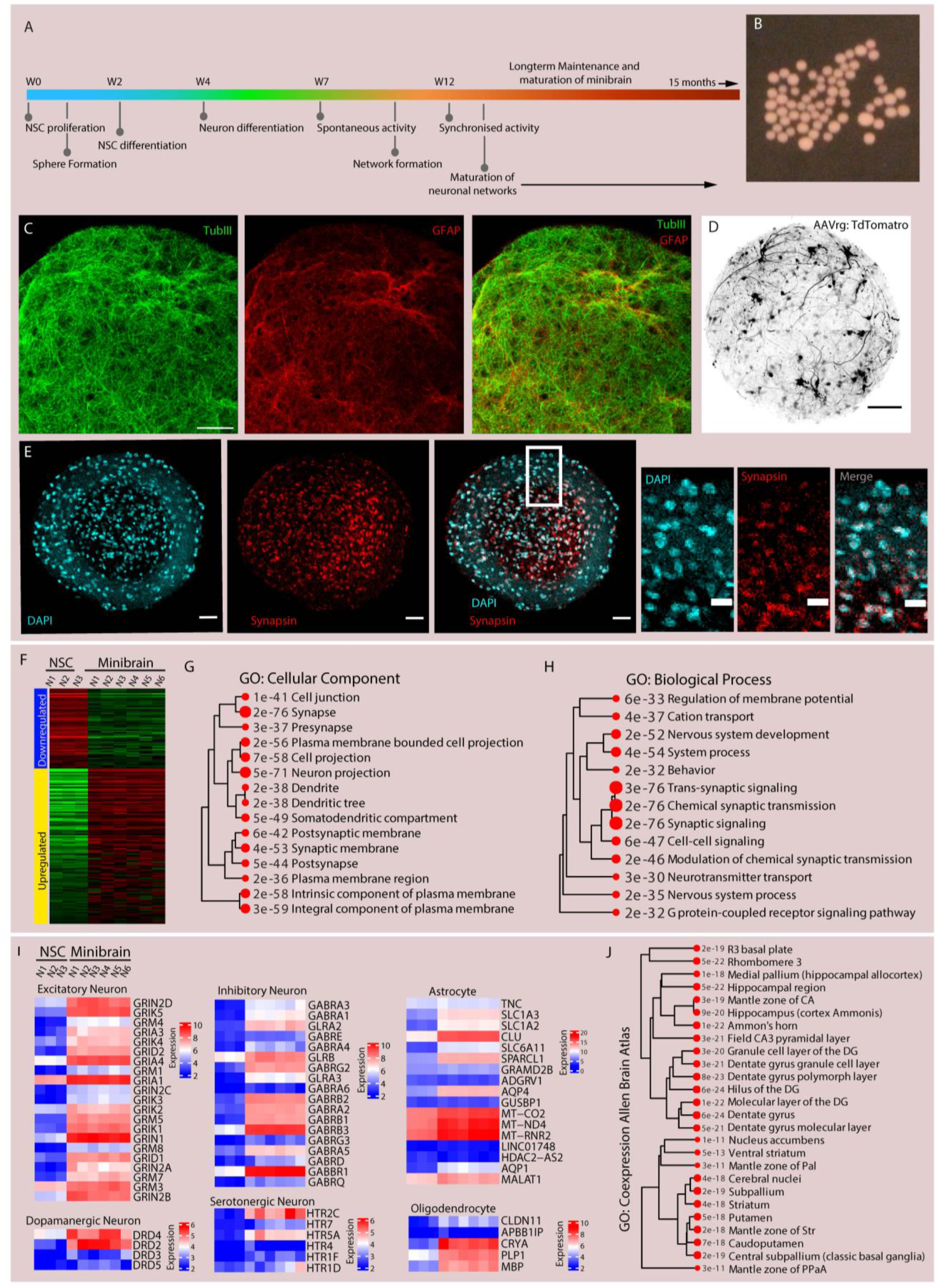
Introduction to minibrain. A) Illustration depicting the development of minibrain adapted from Govindan et al., 2021. B) Photograph of minibrains in a 6 well plate. C) 7-month-old minibrain display expression of beta-TubIII (green) and GFAP (red), scale: 50 μm. D) AAVrg virus mediated tdTomato labelled neurons in minibrain display elaborate projection (black). E) In 3-month-old minibrain, many cells in minibrain express synapsin, a synaptic marker (red), evident by colocalization of some DAPI (blue) cells. Scale: 100 μm. (right) Zoom in of images on the left scale: 20 μm. F) Heat map of differentially enriched genes in NSCh^IPSC^ and 4-month-old minibrains. G) & H) Gene ontology enrichment of cellular components and biological processes in minibrain enriched transcriptome. I) Heatmap of genes enriched in excitatory neuron, dopaminergic neuron, inhibitory neuron, serotonergic neuron, astrocytes and oligodendrocytes. J) Gene ontology enrichment of “Co.expression allen brain atlas gene set” in the minibrain transcriptome.

**Figure 2:**
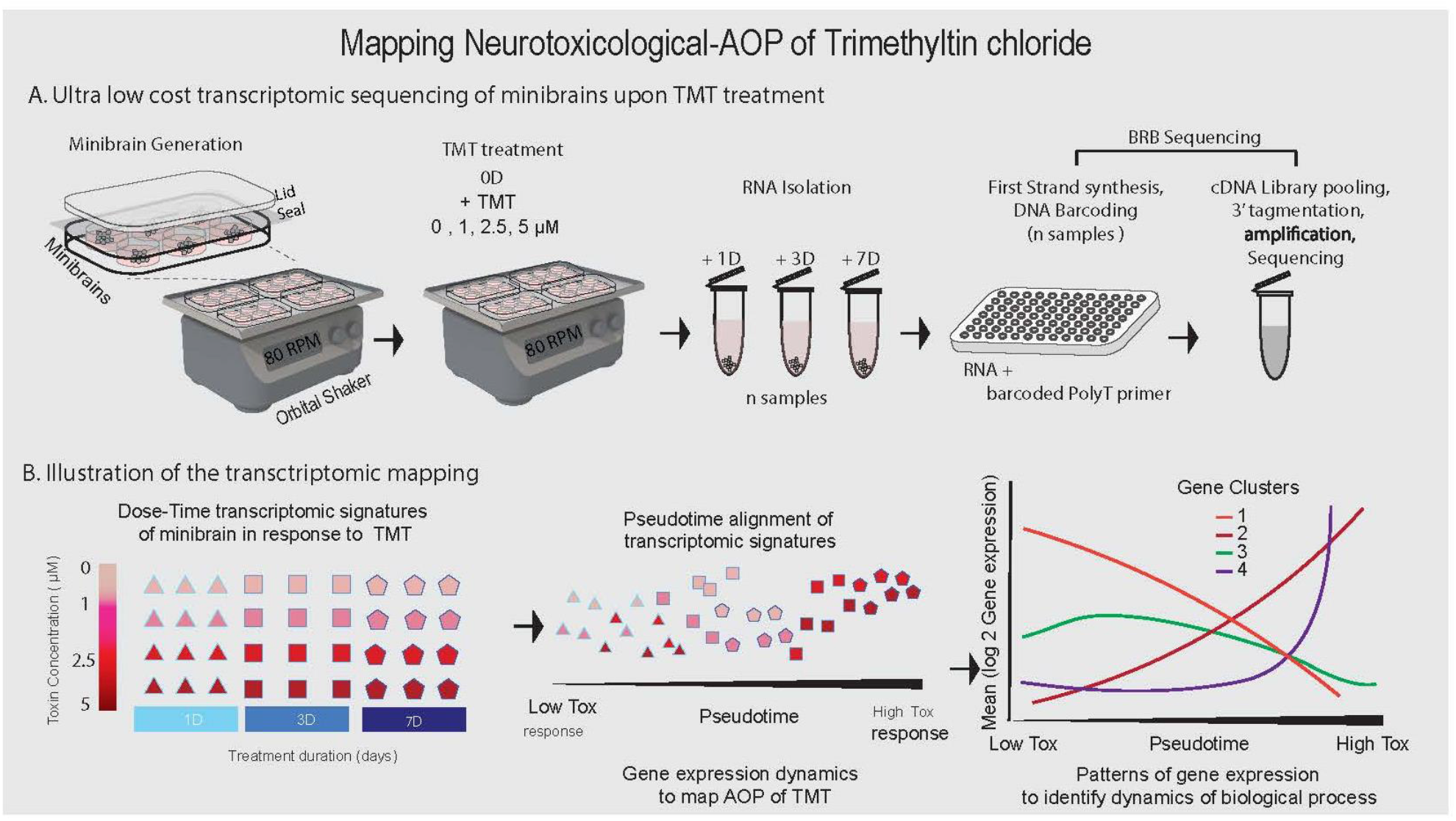
RNA sequencing pipeline using minibrains and BRB sequencing tor toxicity study. A) Pipeline illustrating minibrain generation, dose time treatment of TMT followed by BRB-seq based library preparation and next generation sequencing. B) Analysis pipeline illustrates utilizing the transcriptomic responses to TMT dose-time treatment for identifying gene expression patterns that will be used to identify adverse outcome events of TMT induced neurotoxicity.

**Figure 3:**
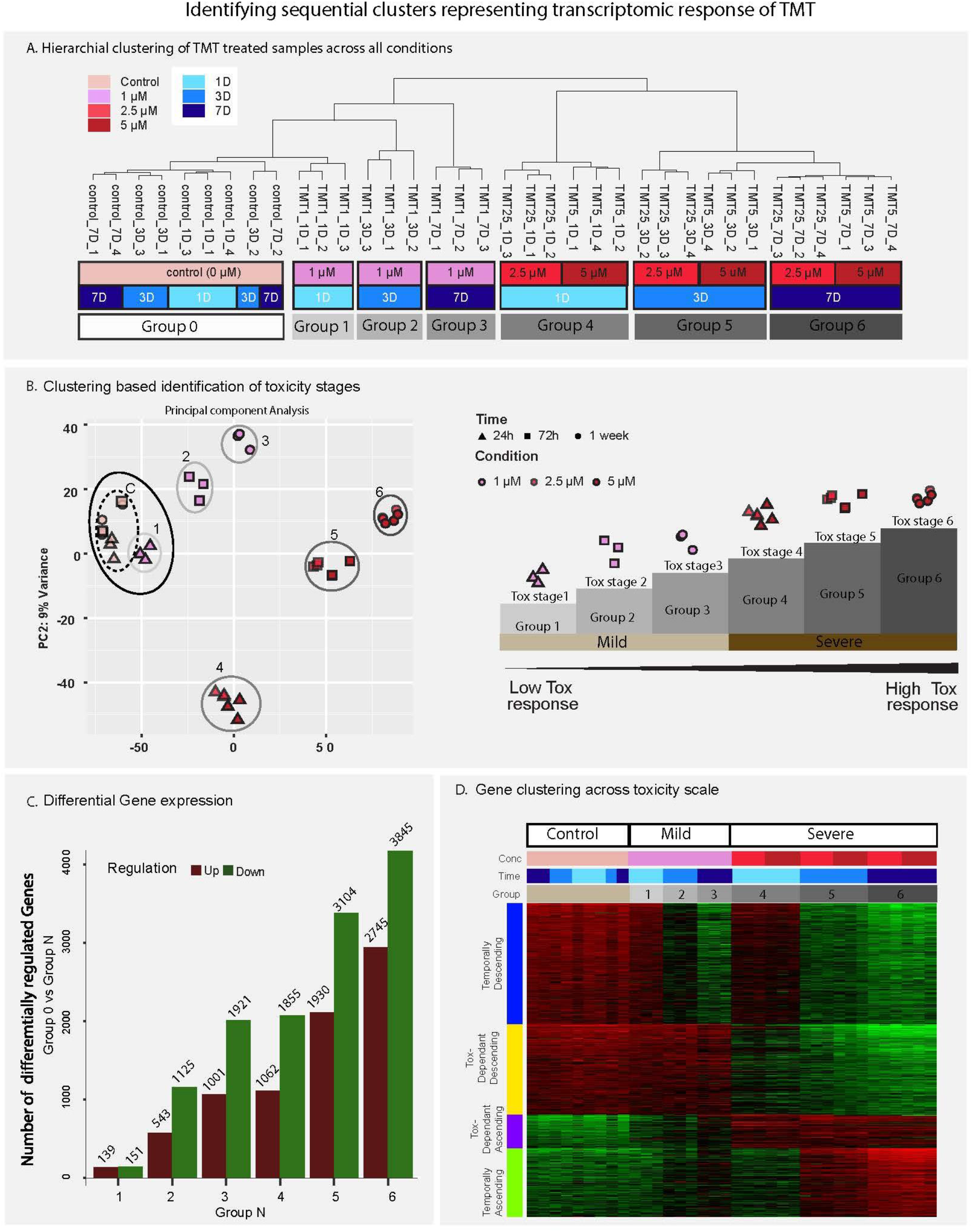
Identifying progressive toxicity based on transcriptomic signature. A) Hierarchical clustering of TMT’s dose time transcriptomic signature. B) Principal component analysis of TMT’s dose time transcriptomic signature (left). Progressive groups were ordered based on the hierarchical clustering and principal component to create a neurotoxicity axis representing low transcriptional response to high transcriptional response (B, right). Samples that clustered together with control in A was named mild and samples that clustered separately in A was named severe groups (B, right). C) Differential enrichment analysis of progressive groups versus control showed progressively increasing transcriptomic response (i.e., differentially enriched genes) of minibrains along the neurotoxicity axis. D) Gene expression clustering of most variable genes reveal time dependent and toxicity severity dependent (Tox-dependent) upregulation and downregulation of genes.

**Figure 4:**
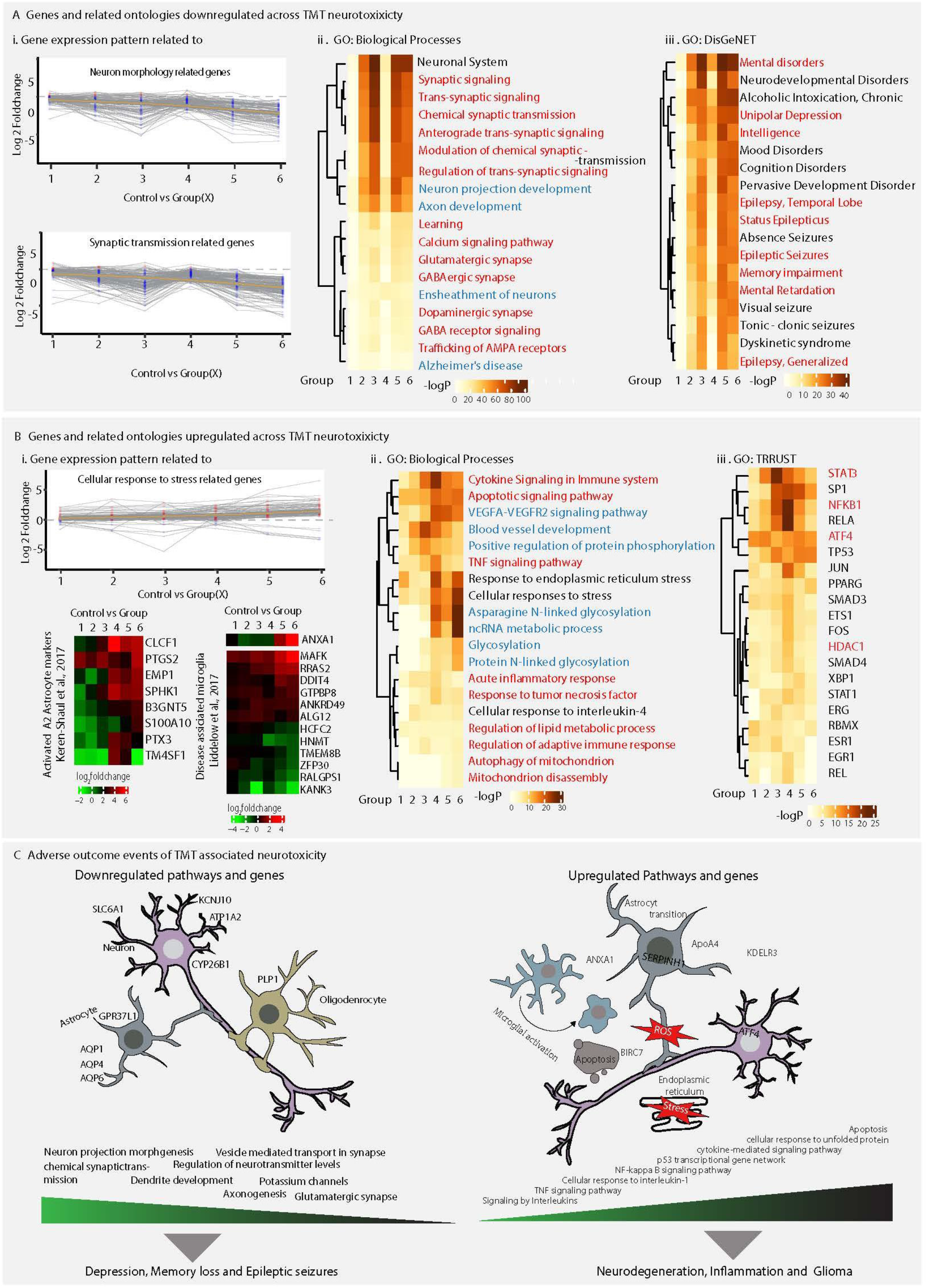
Select adverse outcome events during TMT induced neuro toxicity that leads to pathogenesis. A) Downregulated genes and related ontologies across neurotoxicity axis (I) gene expression pattern of neuron morphology and synapse related genes. Gene ontology analysis for (II) biological processes and (III) DisGenNET for downregulated genes. B) Upregulated genes and related ontologies across neurotoxicity axis. (I, top) Gene expression pattern of cellular stress related genes. (I, bottom) heatmap of activated A2 astrocytes and disease associated microglia. Gene ontology analysis for (I) biological processes and (II) TRRUST for upregulated gene. Gene ontology terms in A, B are highlighted in red if already reported and in blue if notable and unreported in previous TMT neurotoxicity studies. C) Illustration representing various adverse events associated with upregulated and downregulated genes across neurotoxicity axis.

**Figure 5:**
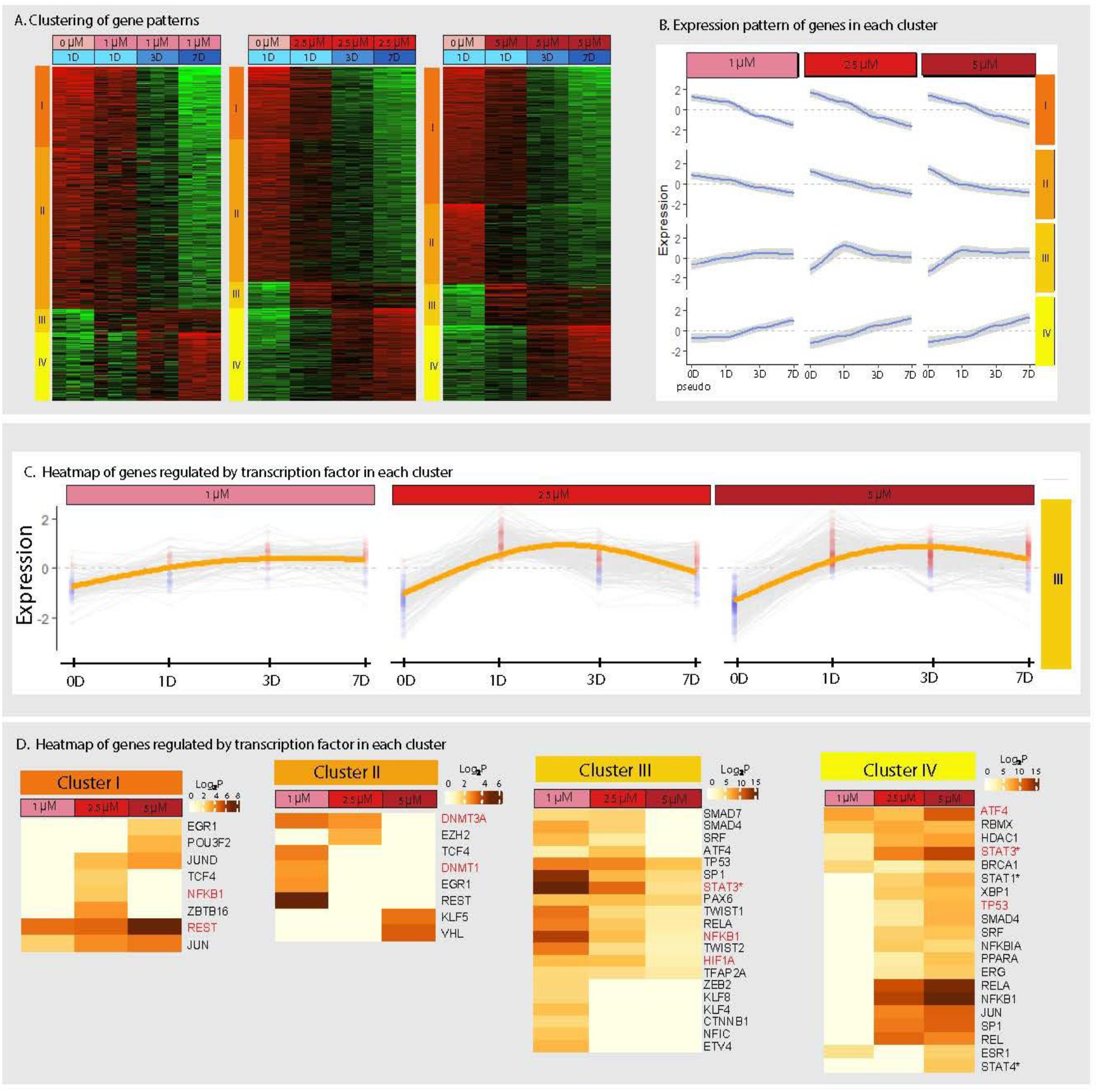
Identifying distinct gene patterns contributing to TMT Neurotoxicity. A) Clustering of genes was performed by K mean clustering algorithm to identify 4 distinct patterns during TMT exposure. 0 μM 1-day exposure was considered as no exposure control (Pseudo 0D). Changes in gene expression across time was used to cluster genes within each dose. B) Expression pattern of genes in each cluster in Cluster I showed drastic downregulation of genes, Cluster II showed progressive downregulation, Cluster III showed upregulation of genes as early as after 1-day(acute) exposure and Cluster IV showed slower and consistent upregulation reaching the maximum at 7-day (chronic) exposure. The graph is a smoothened line plot that shows average expression of genes in blue, grey area represents the maximum and minimum gene expression. C) Cluster III specifically consists of genes that are highly upregulated as early as after 1-day (acute) TMT exposure which is attenuated in consecutive exposures. These lists of genes can serve as markers of early detection of neurotoxicological stress. Graph represent expression (Kmean value) of genes, Red dots indicate gene expression value above 0, Blue dot indicates gene expression value below 0 at a one timepoint for one gene. D) Heatmap representing enrichment for transcription factor governed gene regulatory networks in each cluster. Transcription factors known to be previously associated with TMT induced neurotoxicity is highlighted in red.

### BRB-seq library sequencing and data processing

The libraries were sequenced on Illumina NextSeq 500 platform as previously described at a depth of 3 – 4 million raw reads per sample (Alpern et al., 2019).

### Bioinformatic analysis

The sequencing reads were demultiplexed using the BRB-seq Tools suite (Alpern et al., 2019) and were aligned against human genome (hg38) using STAR (Dobin et al., 2013) and HTSeq (version 0.9.1) (Love et al., 2014) for producing the count matrices. Gene expression analysis was performed using iDEP platform, genes with at least 0.5 counts per million in at least 4 samples were chosen for analysis for figure 1 and genes with at least 0.5 counts per million in at least 9 samples were chosen for analysis for figure 3–5. Hierarchical clustering was performed using 2000 most variable genes centered by subtraction of mean and division of standard deviation. Principal component analysis was performed in the iDEP platform. K mean clustering of genes for figure 3 was performed on 2000 most variable genes across all samples. K mean clustering of genes for figure 5 was performed on 1200 most variable genes of the low condition samples, and 2000 most variable genes of the moderate and high condition samples. Genes were normalized by the mean center for K mean clustering analysis. For differential enrichment analysis, a false discovery rate of 0.1 and a minimum fold change of 2 was given as parameters for DESeq2 analysis in the iDEP platform. Gene ontology analysis was performed in iDEP platform for figure 1. Metascape was used to perform gene ontology analysis using the parameters of the express analysis for figure 4 and 5. Plot generations and further analyses was performed using R version R 4.0.2.

## Results

Minibrain is a human 3D brain *in vitro* model developed in our lab that is composed of complex neuronal and glial cellular subtypes modelling various cellular interactions that happen in the human brain within micrometer scales (Figure 1 A-C) (Govindan et al., 2021). Minibrains are generated and differentiated through a slow transition of NSC^hIPSC^ aggregates from proliferation status to neuronal differentiation (Figure 1A)(Govindan et al., 2021). Minibrains can with ease/cost effectiveness be mass generated/maintained in suspension using culture plates, agitators and a cell culture incubator (Figure 2A)(Govindan et al., 2021). The small size of the minibrain i.e., 0.5-0.6 mm on an average allows them to be amenable to multiplexing, upscaling various experimental conditions for mass scale chemical screening and disease modelling studies (Govindan et al., 2021). Minibrain showed immunoreactivity to beta-Tubulin III and GFAP antibodies confirming the presence and even distribution of neurons and glia respectively (Figure 1 C). 4-month-old minibrains contain differentiated neurons that display synaptic markers and elaborate neuronal morphology (Figure 1 D, E).

To better characterize the minibrain model system we performed transcriptomic analysis of non-differentiated NSC^hIPSC^ and minibrains (3 month old) generated as described in (Govindan et al., 2021). Genes with increased expression in minibrain samples were associated with neuronal function (neuron differentiation, synapse signaling and morphogenesis) and neuron morphology (cellular components such as axon, dendrite and synapse) GO terms (Figure 1 F-H). Enrichment of genes related to synapse function in minibrain illustrate the presence of active neuronal connections. In our previous study we show action potentials, single neuron and synchronized neuronal network activity within the minibrain through extracellular electrode recordings (Govindan et al., 2021). Genes characteristic of excitatory neurons, inhibitory neurons, dopaminergic neurons, serotonergic neurons, astrocytes and oligodendrocytes were enriched in the minibrain transcriptome (Figure 1 I). Minibrain transcriptome showed enrichment for genes that corresponds to various brain regions such as hippocampus, caudate putamen, striatum, sub pallium, nucleus accumbens and cerebral nuclei of the central nervous system (Figure 1 J). TMT is known to severely affect hippocampal neurons specially and thus the enrichment of hippocampal brain region associated genes in minibrain, makes it a suitable model to study TMT induced neurotoxicity (Homayoon et al., 2015; Pershina et al., 2019).

### Toxicogenomic profiling of TMT induced neurotoxicity

Next, we obtained TMT dose-time treatment associated minibrain transcriptome through BRB-seq of minibrains treated with 0 μM (control), 1 μM (low dose), 2.5 μM (moderate dose) and 5 μM (high dose) of TMT for 1-day (Acute), 3-day (intermediate) and 1-week (chronic) duration (Figure 2, Supplementary Figure 1). We analyzed 17,527 genes across the dose-time transcriptional response to identify the molecular events during TMT toxicity. To identify gene expression changes along TMT dose-time treatment, we performed differential gene expression analysis of minibrain transcriptomes corresponding to low, moderate, and high dose TMT treatment over acute, intermediate, and chronic exposure in comparison to acute control non treated samples (0 μM, 1 day) (Supplementary Figure 1 - 3). At each exposure time points, the number of genes differentially expressed in comparison to control increased based on the dose of treatment i.e., low < moderate < high (Supplementary table 1, Supplementary figure 2,3). However, the ratio of number of differentially enriched genes of low dose to high dose treatment (acute: 0.08, intermediate: 0.28, chronic: 0.43) was much lower than the ratio of number of differentially enriched genes of moderate dose to high dose at each exposure time points (acute:0.90, intermediate:0.89, chronic: 1.01) (Supplementary table 1). This illustrates that there is a saturation of the neurotoxic effect on the transcriptome towards the moderate and high dose conditions.

### Toxicity severity based transcriptional divergence map of TMT toxicity

To identify the transcriptomic-response divergence of minibrains upon exposure to TMT toxicity, we performed hierarchical clustering and principal component analysis of minibrain transcriptomes of all dose time combinations along with control minibrain transcriptomes. The analysis revealed 2 major clusters with multiple subclusters (Figure 3A). To correlate transcriptional response based on the severity of TMT neurotoxicity, we ranked and grouped the sample clusters based on their transcriptional divergence from the control samples. All control samples clustered together which was termed as Group control, the rest of the samples clustered themselves progressively away based on the severity of the treatment (Figure 3A, B). Low dose-acute treatment samples and Group control clustered closest in hierarchical clustering but existed as separate subclusters both in PCA and Hierarchical clustering (Figure 3 A, B). Low dose-acute treatment sample cluster represented the earliest and mildest transcriptional response upon exposure to TMT and was termed as Group 1. Next, clusters containing low dose -intermediate and -chronic treatment samples were termed as Group 2 and Group 3 respectively which were situated progressively farther away from Group 1 but still clustered together with Group 1 and Group control. Moderate and high dose samples clustered together reflecting a saturation of the transcriptional response at these TMT doses (Figure 3A, B). Moderate and high dose samples resolved into subclusters representing -acute, -intermediate and -chronic treatment which were termed as Group 4, 5 and 6 respectively. Group 1, 2 and 3 clustered together with Group control representing a milder transcriptional divergence from control samples and hence were termed as mild toxicity group (Figure 3A, B). Group 4, 5 and 6 showed severe transcriptional divergence from Group control and hence were termed as severe toxicity group (Figure 3 A, B). This illustrates that the transcriptional response of minibrains correlates with the severity of the TMT treatment, i.e., minibrain treated with severe doses of TMT display higher transcriptional divergence compared to minibrains treated with milder doses (Figure 3B). Next, through a series of analysis we validate that the transcriptional divergence/response of minibrain across groups to recapitulate and identify known and unknown molecular pathways and mechanism involved in TMT induced neurotoxicity.

We aligned Group 1 to Group 6 on an axis terming it as the neurotoxicity axis to reflect the severity of the transcriptional response upon exposure to TMT (i.e., from mild transcriptional divergence to severe transcriptional divergence). Next we mapped the gene expression changes in each group’s transcriptome in comparison to the control transcriptomes (Figure 3C, Supplementary table 3). Differential gene enrichment (DEG) analysis of each group in comparison to control revealed progressively higher number of DEG genes across the order of grouping i.e., Group 1 to Group 6 reflecting the severity of the toxicity effect (Figure 3 C). We observed higher number of downregulated genes compared to upregulated genes in all severity groups (Supplementary table 3, Figure 3 C).

Majority of genes related to neuron morphology and neuronal synapse were downregulated across groups (Figure 4 A (i)). Notable significant gene ontology terms enriched for downregulated genes was associated with synapse function, axon development, glutamatergic synapse, GABAergic synapse, dopaminergic synapse, calcium pathway, trafficking of AMPA receptors and ensheathment of neurons, learning and Alzheimer’s disease (Figure 4 A (ii)). Many genes were progressively and more significantly downregulated in the severe effect group than the mild effect group which included i) *PLP1*, a myelin component produced by oligodendrocyte required for neuron myelination and ii) *POU3F2*, a neuronal transcription factor which showed immediate downregulation (Figure 4 C, Supplementary table 4). Gene ontology enrichment through DisGeNET database (a curation of gene sets related with various disease) on downregulated genes revealed disorders related to mood, epileptic seizures and learning (supplementary table 5, Figure 4 A, C). Gene ontology analysis of downregulated genes across the neurotoxicity axis recapitulated previously described pathomechanisms such as i. demyelination, ii. learning impairment, iii. loss of glutamatergic synapses, iv. loss of neuronal function, v. epilepsy and vi. calcium homeostasis dysregulation involved in the neuronal dysfunction associated with TMT neurotoxicity (Supplementary table 4,5, Figure 4C) (Veronesi et al., 1991; Feldman et al., 1993; Halladay et al., 2006; Piacentini et al., 2008; Pershina et al., 2019).

Indicative of neuronal stress upon TMT exposure we observed an upregulation of genes related to cellular stress (Figure 4 B(i)). In concurrence with previous studies upregulated genes in TMT treated samples were associated with cytokine signaling, apoptotic pathway, mitochondrial dysfunction, altered lipid metabolic processes and TNF pathway (Figure 4B (ii), supplementary table 4) (Figiel and Dzwonek, 2007; Misiti et al., 2008; Lattanzi et al., 2013; Ferraz da Silva et al., 2018). The TMT induced minibrain transcriptome was enriched with activated astrocyte and disease associated microglial gene signatures (Figure 4 B (i), 4C) (Keren-Shaul et al., 2017; Liddelow et al., 2017). We observed enrichment for GO terms VEGFA -VEGFR2 signaling pathway, and blood vessel development related to angiogenesis which is indicative of astrocytes upregulating endothelial genes, a mechanism that is observed in astrocytes upon stress or in a tumorous environment (Figure 4 B, C) (Brumm et al., 2017). *ANXA1*, which is known to be involved in microglial cells activation required for phagocytosis was upregulated in severe -intermediate and -chronic exposure groups suggesting microglia activation (Figure 4B (i, bottom right), 4C). Previous studies in rat and mouse report activation of astrocytes and microglia upon TMT exposure, which is recapitulated in our TMT severity based transcriptomic mapping (Brabeck et al., 2002; Pompili et al., 2011).

We next analyzed the upregulated genes across neurotoxicity axis using TRRUST database (a reference database of human transcriptional regulatory interactions) to identify gene regulatory networks activated in response to TMT toxicity. TMT induced transcripts were enriched for gene regulatory network governed *STAT3, NFKB1, ATF4* transcription factors which were already known to be associated with TMT induced neurotoxicity (Lattanzi et al., 2013) (Figure 4B (iii)). Overall, our neurotoxic genomic assay using minibrain was able to identify biological pathways, molecular events, cellular events and disease pathways activated in response to TMT (Figure 4, Supplementary table 3-5).

Next we clustered genes based on their expression pattern across the neurotoxicity axis which revealed 4 clusters of genes falling under two categories i) toxicity severity dependent and ii) time dependent gene expression pattern (Figure 3E). Toxicity effect dependent gene clusters consisted of genes that showed distinct enrichment between the mild and severe effect groups while time dependent gene clusters were upregulated or downregulated across acute to chronic treatment exposure in both the mild and severe groups (Figure 3E). Since gene clustering across neurotoxicity axis revealed a time course dependent gene expression pattern, we analysed the expression patterns across acute, intermediate and chronic exposures in low, moderate and high TMT doses independently with control dose acute condition (0 μM, 1 day) as the starting point.

Genes clustered based on 4 distinct patterns across acute, intermediate, and chronic exposure in all doses (Figure 5 A, B, Supplementary figure 4). Gene cluster I showed immediate and higher downregulation of genes compared to gene cluster II that showed a steeper downregulation of genes across exposures (Figure 5A, B, Supplementary figure 4). Gene cluster III showed an immediate and higher upregulation of genes at acute exposure while gene cluster IV showed a progressively increasing upregulation of genes across exposures (Figure 5 A, B, Supplementary figure 4). We performed TRRUST based ontology analysis on each gene cluster to identify the dynamics of transcriptional gene regulatory networks across our dose-time neurotoxicogenomic TMT assay (Figure 5 D, Supplementary table 6).

### Dynamic gene regulatory modules during TMT induced neurotoxicity

Gene cluster I showed significant enrichment of gene regulatory network associated with *REST*, a transcription factor that regulates gene expression miRNA mediated gene silencing which explains the immediate and higher downregulation across all doses in cluster I (Figure 5 D). *REST* has been previously shown to be activated in ischemic stroke, neuron degeneration and neuronal stress induced down regulation of genes (Hwang and Zukin, 2018; Soga et al., 2021). This implies that miRNA mediated gene silencing is one principal biological event that is immediately activated upon exposure to TMT. Gene cluster I of high dose samples show enrichment for gene regulatory network mediated by *POU3F2*, a transcription factor that regulates excitatory neuron identity and facilitated protective mechanisms against neuronal stress. This suggests that there is an immediate downregulation of neuronal function upon high dose TMT (Figure 5 D) (Pang et al., 2011; Marrocco et al., 2017; Mertens et al., 2018; Herbert et al., 2019). Gene cluster II showed enrichment for gene regulatory network associated with *DNMT1* and *DNMT3A*, key players of DNA methylation dependent gene repression regulation that is known to mediate expression of synaptic plasticity genes in response to stress (Figure 5 D) (Feng et al., 2010; Moore et al., 2013; Bayraktar and Kreutz, 2018; Sun et al., 2020). We were able to detect activation of REST (miRNA) and DNA methylation dependent gene silencing mechanism which were previously established to be involved in TMT neurotoxicity (Imam et al., 2018; Kim et al., 2021).

Genes cluster III shows higher upregulation of genes at acute exposure compared to intermediate and chronic exposure. This suggest that gene cluster III represents **early response genes associated with TMT exposure** whose expression increases phenomenally high during initial stages of TMT toxicity but the upregulation is attenuated during progressive treatment exposures (Figure 5A-C, Supplementary figure 4). Gene cluster III of low, moderate, and high dose treatment samples show enrichment for *TP53, SP1, STAT3, PAX6, TWIST1, RELA, NFKB1, TWIST2, HIF1A* and *TFAP2A* mediated gene regulatory network (Figure 5 D). Notably *RELA, NFKB1, TWIST1, TWIST2, TFAP2A* and *SP1* transcription factors are associated with inflammation, cell death, cancer pathway and metastasis. *TP53* and *STAT3* transcription factor is known to regulate genes that provide protective mechanisms against uncontrolled proliferation, apoptosis and oxidative stress (Aubrey et al., 2016). *PAX6* transcription factor is known to regulate neuroprotective mechanisms, anti-cancerous mechanisms and promote resilience to oxidative stress (Mayes et al., 2006; Ninkovic et al., 2010; Hegge et al., 2018). Enrichment of gene regulatory networks associated with transcription factors that mediate stress, cell death pathways along with neuro protective, anti-apoptotic pathways in cluster III is indicative of a homeostatic mechanism upon TMT exposure. *ZEB2*, a gene nriched in low dose - gene cluster III), causes Mowat-Wilson Syndrome which can both activate and repress gene expression and has been shown to play an important role in activation of reactive astrocytes in response to stress (Vivinetto et al., 2020). Concurrently, we observe increase in genes characteristic of A2 astrocytes known for their neuroprotective nature upon exposure to TMT (Figure 4 B) (Liddelow et al., 2017). Previous studies also indicate such activation of protective astrocytic mechanisms upon initial exposure to TMT treatment in other models (Corvino et al., 2013). Gene cluster IV showed a steady and gradual increase in the upregulation of genes from acute to intermediate to chronic exposures across all doses (Figure 5 A, B). Gene cluster IV showed enrichment for genes regulated by *ATF4, RBMX, HDAC1, STAT3* and *BRCA1* across all doses (Figure 5 D). *RBMX* transcription factor regulatory network is indicative of DNA damage pathways activated progressively during TMT exposures (Matsunaga et al., 2012; Harvey et al., 2017). Low dose samples captured most diverse gene regulatory pathways in gene cluster III i.e., early response gene activation to TMT while high dose and moderate dose samples captured the most diverse gene regulatory pathway in gene cluster IV (Figure 5 D). This illustrates that transcriptome of low dose samples capture the initial and early response events to TMT while moderate and high dose samples capture the severe events involved in TMT induced neurotoxicity. This illustrates that it is crucial to perform dose-time toxicogenomic assays to identify both early to late molecular events in response to a toxicant.

### Distinct STAT pathways are activated based on the dose of TMT treatment

*STAT3* regulated genes are enriched in low, moderate and high dose treatment samples while *STAT1* regulated genes were enriched only in moderate and high dose treatment samples (Figure 5 C, center and right). *STAT1* and *STAT3* transcription factor balance is key in mediating the shift of cell survival to apoptosis and anti-inflammatory to pro-inflammatory pathways (Takagi et al., 2002, 3; Hao et al., 2010, 1; Butturini et al., 2018, 2019, 1). This implies that in severe effect doses (i.e., moderate and high dose samples), the minibrain activates homeostatic mechanism to protect against TMT’s neurotoxicity. *STAT4* regulated genes were enriched specifically in high dose samples (Figure 5C). *STAT4* is a transcription factor downstream of IL12 and IL23 play significant role in demyelination and inflammation activation in the CNS (Pagenstecher et al., 2000; Chitnis et al., 2001; Mo et al., 2008; Anderson et al., 2018; Alakhras and Kaplan, 2021). The specific enrichment of *STAT4* gene regulatory network in the high dose samples indicate that *STAT4* regulated genes can serve as marker for severe stress induced by TMT. We identify here an interplay of distinct STAT signaling pathway based on the severity of TMT dose treatment playing a role in TMT induced neurotoxicity.

## Discussion

TMT exposure has been long reported for causing neurological dysfunction and neurodegeneration in industrial workers dealing with plastic stabilizers (Kreyberg et al., 1992). Due to the lack of human 3D brain models in the previous decade, TMT toxicogenomic studies was performed in rat, mouse and PC12 neurons (Lattanzi et al., 2013). Here we demonstrate the efficacy of a neurotoxicogenomic assay that couples human minibrain model and a highly multiplexed transcriptomics approach (BRB-seq) for toxicity screening. Minibrain transcriptome shows enrichment of genes specific to excitatory (glutamatergic)-, inhibitory- (GABAergic) and modulatory- (dopaminergic and serotonergic) neurons, astrocytes, oligodendrocytes and microglia, brain regions such as hippocampus, striatum, caudate accumbens and putamen. Thus, minibrain could serve as model system to study cellular interactions in the above said brain regions at the molecular level.

TMT toxicity is known to have severe effect on hippocampal region of the brain and the pathomechanism of TMT induced neurotoxicity does not involve impairment of the blood brain barrier making minibrain an ideal model to study TMT induced neurotoxicity (Kreyberg et al., 1992; Little et al., 2002).

The use of transcriptomics for molecular characterization of drug and chemical response is markedly growing in pre-clinical research and pharmaceutical industry. Genome wide expression profiling is a versatile molecular phenotyping approach that offers in-depth analysis of drug mechanism of action and off target effects. The high cost and high labor intensity are principal limitations of the traditional RNA-seq workflows. Recently, several library preparation protocols aiming at reducing the time and cost of RNA-sequencing have emerged ((Bush et al., 2017; Ye et al., 2018; Alpern et al., 2019)). The common aspect of these workflows is the capacity for the early multiplexing of samples achieved via sample tagging during reverse transcription reaction with barcoded oligos. This approach enables to pool up to 384 samples or more in a single tube for subsequent library preparation steps, which in turn makes possible high throughput and high content screening for drug or biomarker discovery research.

Adverse outcome pathway (AOP) is a collection of information that allows us to mechanistically link various events at distinct biological levels in response to chemicals i.e., from ecological to organismal to macromolecular interaction to cellular and molecular level. AOPs allow us to build a knowledge framework to assess the risk of chemical toxicants. Using our dose-time neuro toxicogenomic assay we were able to identify time and severity dependent molecular events, pathways and gene regulatory networks that will contribute to building the AOP for TMT induced neurotoxicity. We identified gene regulatory network indicative of activation of miRNA and DNA methylation based gene silencing mechanism involved in the downregulation of genes upon TMT exposure. We reveal activation of an early molecular homeostatic mechanism upon acute exposure to TMT neurotoxicity that includes anti-apoptotic, pro-apoptotic, neuroprotective and cancer related pathways. Our study recapitulated pathways and genes involved in oxidative stress, microglial activation, astrocyte transition, homeostatic mechanisms, dysfunction of neuron and neurodegeneration that were previously shown to be activated in *in vivo* models of TMT induced neurotoxicity (Kassed et al., 2004; Lattanzi et al., 2007, 2013; Kim et al., 2021). DisGeNET analysis on our TMT induced minibrain transcriptomic data was able to identify previously known TMT induced disease phenotypes such as memory impairment, depression and epilepsy (Fortemps et al., 1978) (Supplementary table 5).

Many therapeutic alternatives have been studied for their protective action against TMT neurotoxicity in pure neuronal cultures and *in vivo* (Shin et al., 2005; Edalatmanesh et al., 2016; Zhao et al., 2016; Hou et al., 2017). Our study presents straightforward toxicogenomic approach as a powerful alternative by building on human minibrain 3D *in vitro* model and low-cost highly multiplexable sequencing. Our toxicogenomic approach is cost efficient amenable for upscaling for screening an array of molecules. Our assay will allow screening candidates for its neuroprotective nature against TMT as well as map the molecular mechanisms involved in the reversal of the pathomechanism. This workflow provides a ready to use platform for high throughput in depth screening that can be translated for the purpose of identifying pathomechanisms of neural disorders and testing potential therapeutics for their activity against neurological conditions.

## Supporting information

Supplementary Figures and legends for Supplementary figures

## 3 Author Contributions

Dr. Subashika Govindan conceived the project, performed experiments, performed analysis and wrote the manuscript. Prof. Adrien Roux supervised the project, performed experiments and contributed to the conception of the project and manuscript preparation. Prof. Luc Stoppini provided critical support in the establishment of the minibrains and TMT neurotoxicity assay. Dr Daniel Alpern and Prof. Bart Deplancke provided support in setting up the neurotoxicogenomic pipeline using BRB-seq and provided critical input to the manuscript.

## 5 Acknowledgments

We acknowledge Laetitia Nikles (HEPIA HES-SO) for technical assistance for minibrain generation, TMT treatment, RNA extraction assistance required for the project.

