## Supplementary Figures and legends for Supplementary figures for "Neuro-toxicogenomic mapping of TMT induced neurotoxicity using human minibrain reveals associated adverse outcome events"

Supplementary Material

### Supplementary Figures

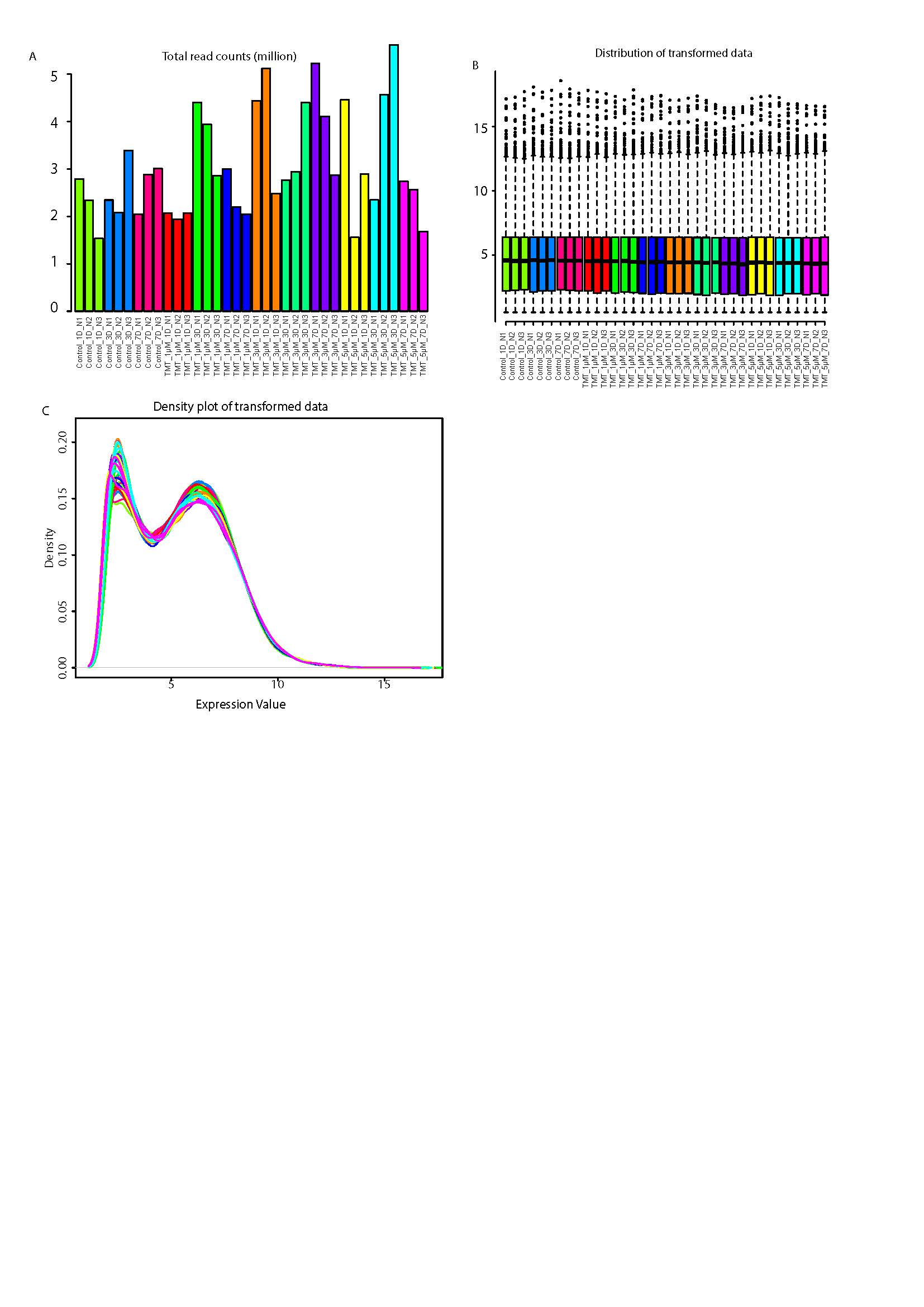

**Supplementary Figure 1.** **Sequencing data distribution across all samples of neurotoxicogenomic assay**. A) Shows the number of reads obtained per sample. B) shows distribution of transformed gene expression of all genes in each sample. C) Shows gene expression density of all genes in each sample. A, B and C shows similar distribution of gene expression density and distribution across all samples. Data from these samples were used for figure 3-5.

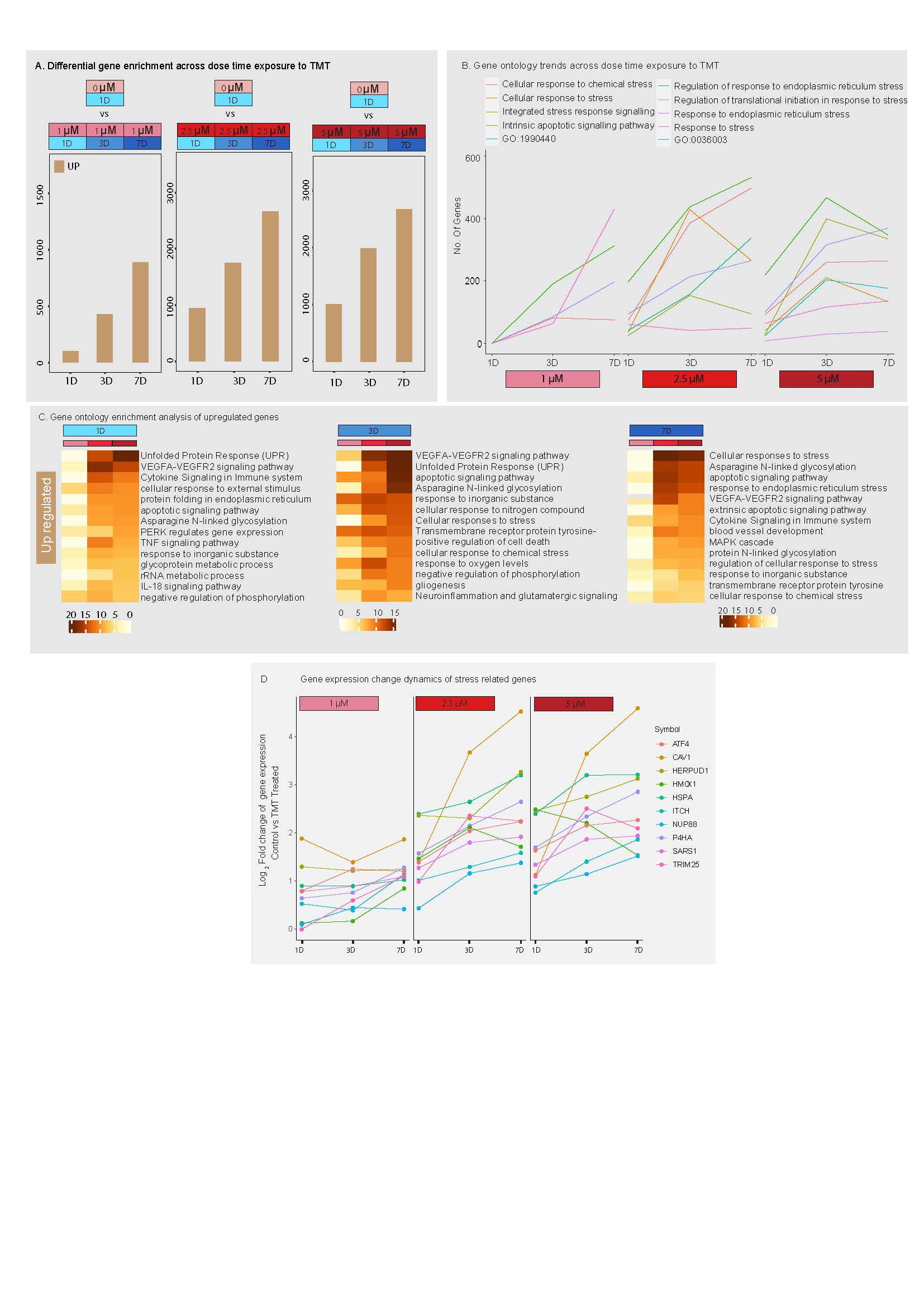

**Supplementary figure 2: Upregulation of genes across dose-time TMT exposure.** A) shows the number of genes upregulated in minibrain upon exposure to distinct doses of TMT across 1-day (1D, acute), 3-day (3D, intermediate) and 7-day (7D, chronic) exposure timepoints (refer to supplementary table 1). B) Graph shows the number of genes enriched in ontology term related to stress across exposure timepoints within distinct doses of TMT. C) shows the enrichment of gene ontology terms of upregulated genes across distinct concentration within each exposure time point. D) shows expression pattern of genes related to stress across exposure timepoints within each TMT dose.

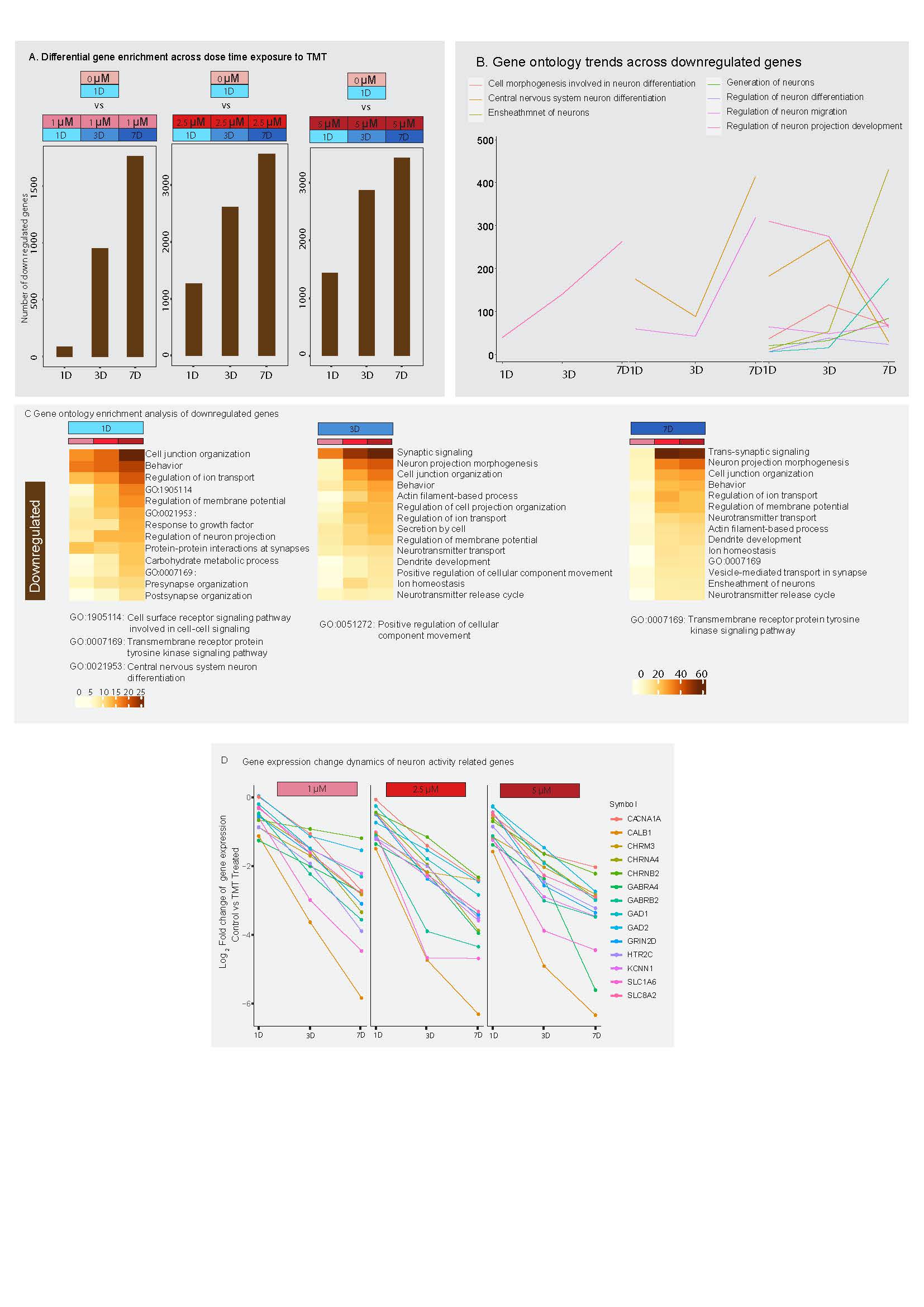

**Supplementary figure 3:** **Downregulation of genes across dose-time TMT exposure.** A) shows the number of genes downregulated in minibrain upon exposure to distinct doses of TMT across 1-day (1D, acute), 3-day (3D, intermediate) and 7-day (7D, chronic) exposure timepoints (refer to supplementary table 1). B) Graph shows the number of genes enriched in ontology term related to neuron function across exposure times within distinct doses of TMT. C) shows the enrichment of gene ontology terms of downregulated genes across distinct doses within each exposure time points. D) shows expression pattern of genes related to neuron activity across exposure timepoints within each TMT dose.

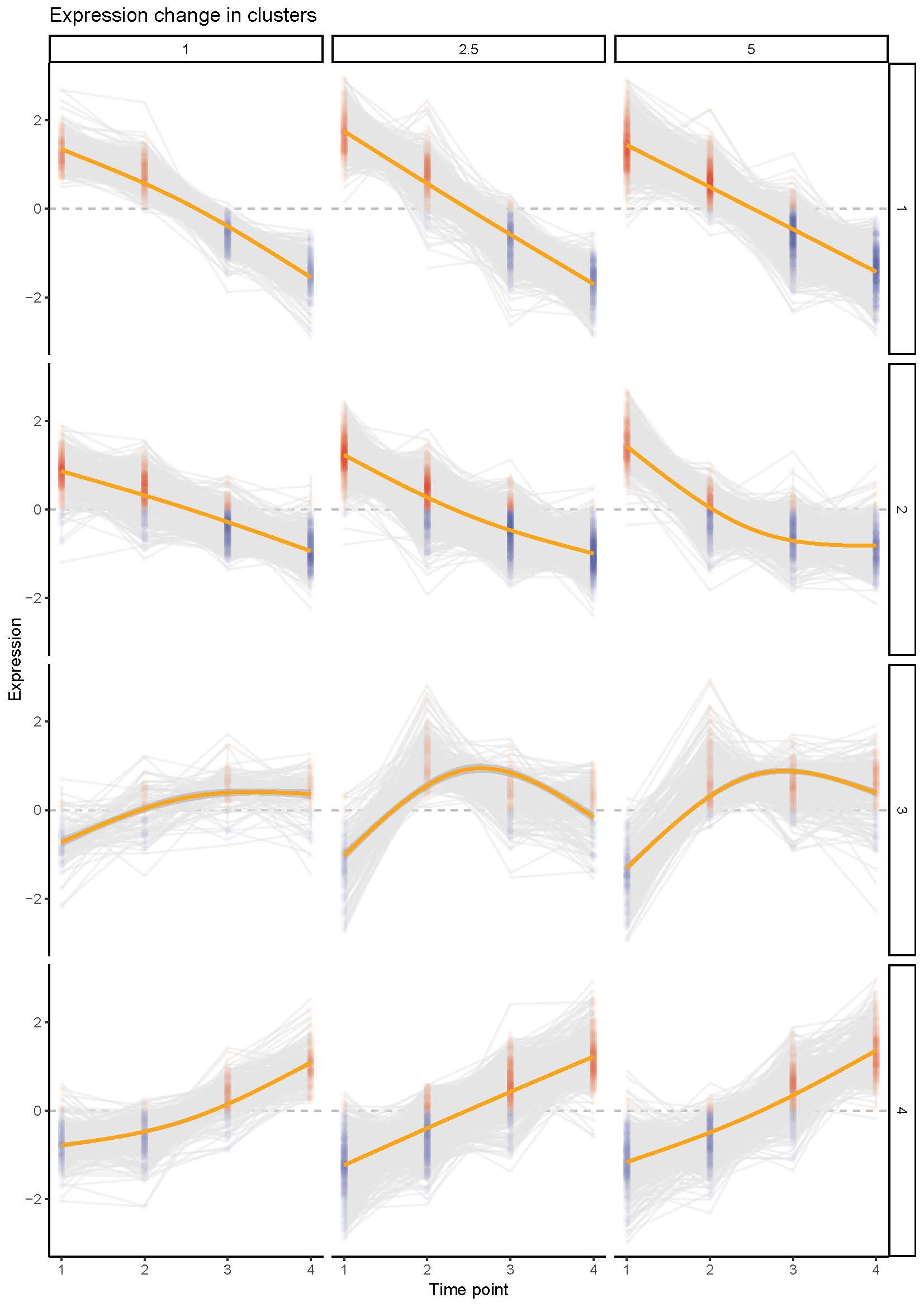

**Supplementary figure 4:** Shows the k-mean value of gene expression across exposure time points within each concentration. Data is related to figure 5. On x axis, 1 = Pseudo OD(0 µM, 1-day), 2 = 1-day, 3 = 3-day and 4 = 7-day exposure time point.

### Supplementary tables

**Supplementary table 1:** List of differentially enriched genes across dose-time exposure of TMT (related to supplementary figure 2, 3).

**Supplementary table 2:** Enrichment of gene ontology terms associated with differentially enriched genes across dose-time exposure of TMT (related to supplementary figure 2, 3).

**Supplementary table 3:** List of differentially enriched genes across groups identified in figure 3.

**Supplementary table 4:** Enrichment of Gene ontology terms associated with differentially enriched genes across groups in figure 4.

**Supplementary table 5:** DisGeNet enrichment associated with differentially enriched genes across groups in figure 4.

**Supplementary table 6:** TRRUST analysis associated gene regulatory network enrichment across gene clusters in figure 5.
